# Molecular and clinical characterization of 5mC regulators in glioma: results of a multicenter study

**DOI:** 10.1101/2020.05.15.098038

**Authors:** Liang Zhao, Jiayue Zhang, Zhiyuan Liu, Yu Wang, Shurui Xuan, Peng Zhao

**Author notes:** Authors contributed equally (Liang Zhao, Jiayue Zhang, Zhiyuan Liu). **Correspondence to:** Peng Zhao, Department of Neurosurgery, The First Affiliated Hospital of Nanjing Medical, University, Nanjing, Jiangsu, 210000, China.

## Abstract

DNA methylation has been widely reported to associate with the progression of glioma. DNA methylation at the 5 position of cytosine (5-methylcytosine, 5mC), which is regulated by 5mC regulators (“writers”, “erasers” and “readers”), is the most critical modification pattern. However, a systematic study on the role of these regulators in glioma is still lacking. In this study, we collected gene expression profiles and corresponding clinical information of gliomas from three independent public datasets. Gene expression of 21 5mC regulators was analyzed and linked to clinicopathological features. A novel molecular classification of glioma was developed using consensus non-negative matrix factorization (CNMF) algorithm, and the tight association with molecular characteristics as well as tumor immune microenvironment was clarified. Sixteen prognostic factors were identified using univariate Cox regression analysis, and a 5mC regulator-based gene signature was further constructed via the least absolute shrinkage and selection operator (LASSO) cox analysis. This risk model was proved as an efficient predictor of overall survival for diffuse glioma, glioblastoma (GBM), and low-grade glioma (LGG) patients in three glioma cohorts. The significant correlation between risk score and intratumoral infiltrated immune cells, as well as immunosuppressive pathways, was found, which explained the difference in clinical outcomes between high and low-risk groups. Finally, a nomogram incorporating the gene signature and other clinicopathological risk factors was established, which might direct clinical decision making. In summary, our work highlights the potential clinical application value of 5mC regulators in prognostic stratification of glioma and their potentialities for developing novel treatment strategies.

## Introduction

Glioma is the most common malignant brain tumor, with an approximate incidence rate of 6/100000 between 2010 to 2014(1). According to the World Health Organization (WHO) classification of central nervous system (CNS) tumors, diffuse gliomas are classified based on histological features(2). Due to the extensive intratumoral heterogeneity and differences of molecular characteristics, the prognosis of glioma patients diverse a lot. Debulking surgery combined with chemo- and radiotherapy has become the standard therapeutic option for glioma (3). However, glioma cells have the ability to diffusely infiltrate into normal brain tissues, which make it hard to achieve complete resection and even cause tumor recurrence. Therefore, it is urgent to explore the potential mechanisms of the development and progression of glioma.

DNA methylation is an essential regulator of epigenetic modification that can affect gene expression without changing DNA sequences (4). This change takes place in various cellular processes during mammalian development, including transcription, genomic stability, and chromatin structure (5) (6). Growing studies have emphasized the crucial role of DNA methylation when various human diseases occur at the aberrant epigenetic modification, such as cancer (7) (8) (9). Previous studies have reported that DNA methylation is one of the most well-characterized epigenetic changes in glioma (10). DNA methylation aberration on specific gene promoter can affect both clinicopathological and molecular features of glioma, and further, affect the prognosis of patients. For example, O6-methylguanine-DNA-methyltransferase (MGMT) methylation, which is the best-described methylation pattern in glioma, is tightly associated with the DNA alkylating agent, such as the first-line chemotherapeutic drug, temozolomide (TMZ) (11). Methylation of the MGMT promoter has been recognized as a predictive marker for favorable prognosis and sensitivity to TMZ treatment (12). Furthermore, novel glioma molecular subtypes have been established based on DNA methylation profiles and other multi-omics data, and this finding contributed to a better understanding of glioma and may direct personalized treatment (13).

Until now, methylation of cytosine at the fifth carbon position in CpG dinucleotides is regarded as the most important regulation form and has been intensely studied (14). 5-methylcytosine (5mC) is enzymatically regulated by proteins involved in writing, reading, and erasing the modifications (15). The prominent 5mC regulators contain “writers” such as DNA methyltransferase1 (DNMT1), DNMT3A, DNMT3B, “readers” such as methyl-CpG binding domain protein1 (MBD1), MBD2, MBD3, MBD4,methyl-CpG binding protein 2 (MECP2), Nei like DNA glycosylase 1 (NEIL1), Nth like DNA glycosylase 1 (NTHL1), single-strand-selective monofunctional uracil-DNA glycosylase 1 (SMUG1), thymine DNA glycosylase (TDG), ubiquitin-like with PHD and RING finger domains 1 (UHRF1), UHRF2, uracil DNA glycosylase (UNG), zinc finger and BTB domain containing protein 33 (ZBTB33), ZBTB38, ZBTB4 (16) (17) (18). The identification of 5mC regulators accelerated the researches about the underlying regulatory mechanisms of DNA methylation related human diseases (19) (20).

Recently, the indispensable role of 5mC regulators in carcinogenesis has been widely studied. DNMT1 is responsible for maintaining intracellular global methylation and abnormal CpG island methylation in tumors (21). Overexpression of MBD2 keeps the malignant phenotype of glioma by suppressing the activity of BAI1, a tumor suppressor gene (22). Overexpressed CXXC4 and CXXC5 can destroy the stability and function of TET2 by directly interactions in several tumors, such as colorectal and breast cancer (23) (24). Also, novel therapeutic strategies targeting 5mC regulators have been developed, including the DNMT1 antisense oligodeoxynucleotide (25). However, a systematic analysis of the role of 5mC regulators and the complicated associations with both clinical and molecular features in glioma are still lacking.

In our study, we enrolled three independent glioma datasets and comprehensively investigated the expression patterns of 21 5mC regulators based on large-scale transcriptome data. The prognostic values of these genes were identified, and a gene signature based on 5mC regulators was constructed to predict the clinical outcomes of glioma patients. Moreover, glioma samples were classified into two distinct subtypes on the bias of 5mC regulators, and the interaction between this classification and tumor immune was further explored.

## Methods

### Molecular profiles and clinical data for glioma samples

Level 3 gene expression profiles with transcripts per kilobase million (TPM) values based on Illumina HiSeq RNA-Seq platform and clinical information of both GBM and LGG samples from TCGA database were downloaded from the UCSC Xena data portal (http://xena.ucsc.edu/). Briefly, “TCGA TARGET GTEx” data hub of the Xena platform, which recompute the RNA-seq data from TCGA, Therapeutically Applicable Research To Generate Effective Treatments (TARGET) and Genotype Tissue Expression Project (GTEx) database to create consistent metadata free of computational batch effects, was enrolled to compensate for insufficient normal brain samples in TCGA dataset. In this dataset, we filtered LGG and GBM samples according to the TCGA barcode and selected normal brain samples in the GTEx cohort (cortex, frontal cortex (Ba9), and anterior cingulate cortex (Ba24)) using the “UCSCXenaTools” R package (26). Also, RNA-seq data with raw counts of glioma samples in the TCGA database were downloaded for differential gene expression analysis. Gene expression data with fragments per kilobase million (FPKM) values based on Illumina HiSeq platform of 182 LGG and 139 GBM specimens were obtained from the CGGA website (http://www.cgga.org.cn/) (up to May 6, 2020). The REMBRANDT brain cancer dataset comprising Affymetrix U1332 Plus microarray gene expression data for samples from 671 patients were downloaded from the Gene Expression Omnibus database (GSE108474). Patients without clinical and pathological information were removed from the REMBRANDT cohort. The somatic mutation and copy number variation (CNV) profiles of glioma samples in the TCGA cohort were downloaded from the cBioPortal website (https://www.cbioportal.org/). The TCGA and CGGA dataset were defined as discovery and validation cohort, and the REMBRANDT cohort was used for further evaluating the prognostic value of our risk model mentioned below. Our study fully meets the TCGA publication requirements (http://cancergenome.nih.gov/publications/publicationguidelines).

For the CGGA dataset, RNA-seq data with FPKM values were transformed into TPM format values, which are more suitable for comparing the gene expression levels between different samples. Raw data from the REMBRANDT cohort were pre-processed using the RMA function for background adjustment in the “Affy” R package (27) and then normalized using the “limma” package (28). Gene symbols corresponding to the probe ID in the REMBRANDT dataset were annotated using the “hgu133plus2” R package. The average gene expression value was adopted when multiple probe IDs match a single gene symbol.

In the TCGA cohort, molecular subtype information of gliomas was obtained from published literature, including Verhaak 2010 classification(29), Wang 2017 GBM classification(30), and LGG molecular classes(31). The classification data of gliomas in the CGGA cohort were downloaded with the clinical file.

### Gene list of 5-methylcytosine (5mC) DNA methylation regulators

From a published research (32), we collected a total of 21 5mC regulators. Regulators were classified into three distinct types according to their biological functions, including the “writers” (DNMT1, DNMT3A, DNMT3B), “readers” (MBD1, MBD2, MBD3, MBD4, MECP2, NEIL1, NTHL1, SMUG1, TDG, UHRF1, UHRF2, UNG, ZBTB33, ZBTB38, ZBTB4), and “erasers” (TET1, TET2, TET3). Then, we confirmed that all these genes were available in TCGA and CGGA datasets, while the expression data of NEIL1 was not included in the REMBRANDT cohort. Thus, the REMBRANDT dataset was excluded for exploring the expression profiles of 5mC regulators in glioma.

### Consensus Non-negative matrix factorization

Based on 21 5mC regulators, a clustering method, consensus non-negative matrix factorization (CNMF), which is an effective dimension reduction algorithm for identifying molecular patterns from high-dimensional data, was applied to perform classification of all glioma samples in the TCGA cohort. CNMF method was executed using the “ExecuteCNMF” function implanted in the “CancerSubtypes” R package (33). To guarantee the stability and robustness of the clustering result, we set the detailed parameters during this process as follows: clusterNum = 2 to 4, nrun = 30. The consensus heatmap was used to evaluate the performance of the clustering. Also, silhouette width, which is an index measuring the similarity of individual objects mapping to its identified cluster compared to neighboring clusters, was plotted to assess the quality of the classification. Furthermore, PCA analysis was carried out to present the gene expression patterns in different groups. The distribution of clinicopathological factors in the identified subtypes was investigated, including WHO grade, IDH mutation status, MGMT methylation status, 1p19q deletion, and TCGA Verhaak classification.

### Functional enrichment analyses

Gene set enrichment analysis (GSEA) was used to determine how the signaling pathways differ among two glioma clusters in the TCGA cohort. Firstly, the differentially expressed genes between clusters were identified using the “DESeq2” R package (34). Next, the log2 Fold-Change values of differential gene expression from high to low were used as input to align all glioma samples. GSEA gene sets, including Gene Ontology (GO) (C2), Gene Ontology (GO) (C5), and hallmark (H) gene sets were downloaded from the Molecular Signatures Database (v7.0) (https://www.gsea-msigdb.org/gsea/msigdb/index.jsp). Gene sets related to proneural and mesenchymal GBM features were derived from previous research(35). GSEA was carried out with 10000 times permutations using the “fgsea” R package (36). P-values were adjusted using the discovery rate (FDR) method, and an FDR value less than 0.05 was considered as statistically significant.

GO and the Kyoto Encyclopedia of Genes and Genomes (KEGG) analyses were performed to annotate the functions of the genes we interested. Genes with log2 Fold-Change value >1 and adjusted P-values <0.05 were filtered and submitted to the Metascape website (https://metascape.org/), which is an online tool for gene annotation and analysis resource. GO biological process, KEGG pathways, Reactome gene sets, and canonical pathways were selected during this process. Gene terms with P-value <0.05 were regarded as statistically significant.

### Identification of prognostic 5mC regulators and construction of a 5mC regulator-based risk model

A total of 683 glioma samples, including 518 LGG and 165 GBM patients, with both survival information and gene expression data were filtered. Then, overall survival-related 5mC regulators were identified based on the univariate Cox regression analysis, and genes with P-values <0.05 were selected for the next studies. Hazard ratio (HR) and 95% confidence interval (CI) were calculated and visualized using the “ezcox” R package (37). Notably, the least absolute shrinkage and selection operator (LASSO) with L1-penalty, a well-established linear model for selecting optimal genes with strong prognostic values from high-dimensional data(38), was performed in the present study. Based on the survival-related 5mC regulators in the TCGA cohort, which were filtered by the univariate Cox regression analysis, the number of key genes was narrowed down by the LASSO method. In this method, regression coefficient estimates of unimportant factors can be shrunk to zero; therefore, a number of candidate genes were eliminated. Ten-fold cross-validation was used to determine the optimal value of λ(39), and we chose λ based on minimum criteria. Here, we subsampled the dataset 1000 times and chose the 5mC regulators, which recurred more than 500 times (39). The “glmnet” package was used to perform a LASSO regression analysis.

Genes with the most significant prognostic values were further subjected to the multivariate Cox regression analysis. The risk model based on the 5mC regulator signature was constructed by combining the gene expression levels and the corresponding regression coefficients that were derived from the multivariate Cox regression analysis. The final calculation formula for each glioma patient was as follows:

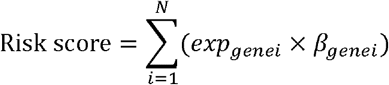

N in the formula represents the number of genes in this risk model, exp_genei_ and β_genei_ mean the expression level and the regression coefficient of a specific gene, respectively. Patients were separated into a high and low-risk group according to the optimal cutoff using the “surv_cutpoint” function in the “survminer” R package. The overall survival differences between two risk level groups were compared using Kaplan-Meier survival analysis based on the log-rank test. To evaluate whether this 5mC regulator-based signature can serve as an independent prognostic biomarker, HR and 95% CI of this signature were calculated from the univariate and multivariate Cox regression analyses. In these methods, other clinical-relevant covariates were included if this detailed information were available, such as age at diagnosis, gender, Karnofsky Performance Scale (KPS), IDH mutation status, MGMT methylation status, etc. Time-dependent receiver operating characteristic (ROC) curves were plotted to investigate the performance of this gene signature in predicting the clinical outcomes of patients at different time points.

### Estimation of immune cell infiltration in tumor samples

Single-sample GSEA (ssGSEA) can calculate the enrichment score of a particular gene set for each sample, and the normalized enrichment score often represents the degree of gene expression ranks inside and outside the gene set. Here, the proportions of immune cells infiltrated in tumor samples were quantified using the ssGSEA method implemented in the “GSVA” R package (https://bioconductor.org/packages/release/bioc/html/GSVA.html). Curated marker gene sets of twenty-four types of immune cells were obtained from a published study (40), and further used as the input file for ssGSEA. Among the gene list, innate immune system immune cells, including neutrophils, mast cells, eosinophils, natural killer (NK) cells, NK CD56dim cells, NK CD56bright cells, dendritic cells (DCs), immature DCs (iDCs), activated DCs (aDCs), and macrophages, as well as adaptive immune cells, including B, T central memory (Tcm), CD8+ T, T effector memory (Tem), T follicular helper (Tfh), gamma delta T (Tgd), Th1, Th2, Th17, and regulatory T (Treg) cells, were included. The output file containing the enrichment score of each kind of immune cell in a single sample was obtained and further used to visualize the discrepancies of immune infiltration in different groups.

### Estimation of tumor purity and enrichment of immune-related signaling pathways

ESTIMATE is a widely used method to speculate the abundance of immune and stromal cells in the malignant tumor samples using gene expression profiles (41). Following the instruction manual of “ESTIMATE” R package, we calculated the enrichment scores of both immune and stromal cells in glioma samples and tumor purity, which indicates the fraction of malignant cells in the tumor niches, were also obtained.

Manually curated gene list of immunosuppressive signaling pathways was used to further characterize the role of 5mC regulator-based signature in regulating pro-tumor activities. These gene sets comprised of activated stroma (42) and several glioma-related immunosuppressive factors (43), including immunosuppressive cytokines and checkpoints, tumor-supportive macrophage chemotactic and skewing molecules, immunosuppressive signaling pathways, and immunosuppressors. Finally, we used ssGSEA to qualify the enrichment score of these pathways in glioma samples.

### Construction and evaluation of a nomogram

The nomogram was generated to quantify the likelihood to survive in 1, 2, 3, and 5-year for glioma patients. Multivariable Cox regression analysis was applied with the clinicopathological factors, such as gender, age, MGMT methylation status, KPS, IDH mutation status, risk score, etc. Detailed covariates in this process were adopted according to the available information in the TCGA or CGGA cohort. The nomogram was plotted using the “rms” package. The calibration plot is a credible method to check whether the nomogram fits on randomized patients, and therefore evaluate the performance of this developed model. Meanwhile, the concordance index (C-index) was used to measure the discrimination of the nomogram in this study.

### Statistical analysis

The LASSO Cox regression analysis was conducted using the “glmnet” R package. Principal component analysis (PCA) analysis was performed and visualized using the “ClassDiscovery” R package. “Survival” and “survminer” R package were used to plot Kaplan-Meier survival curves. The gene interactions among 5mC regulators were obtained from the STRING database (https://string-db.org/), and their correlations were further plotted using the “corrplot” package.

A Chi-square or Fisher’s exact test was performed for checking the differences in sample’s characteristics in different subtypes. A Mann-Whitney U test was used to test the differences in means of continuous data. A two-sided P-value <0.05 was considered as statistically significant for all hypothetical tests. All statistical analyses were performed using R software (version 3.6.3, www.r-project.org).

## Results

### Overview of 5mC regulators in gliomas

Given the vital role of 5mC regulators in the development and progression of human cancers, we firstly investigated the relationships between the expression levels of these genes and clinicopathological features of glioma patients in both TCGA and CGGA datasets. The expression levels of all 5mC regulators were significantly associated with the histology of samples in the TCGA cohort (ANOVA, P <0.001) (Figure 1A). Among them, NEIL1 was highly expressed in normal brain tissues compared with diffuse gliomas, while the rest regulators exhibited the opposite expression patterns. Similarly, we noticed that the expression levels of most of these genes were also related to the WHO grade in the CGGA cohort (ANOVA, P <0.05), except MBD1, NTHL1, and TET3 (Figure 1B).

**Figure 1.**
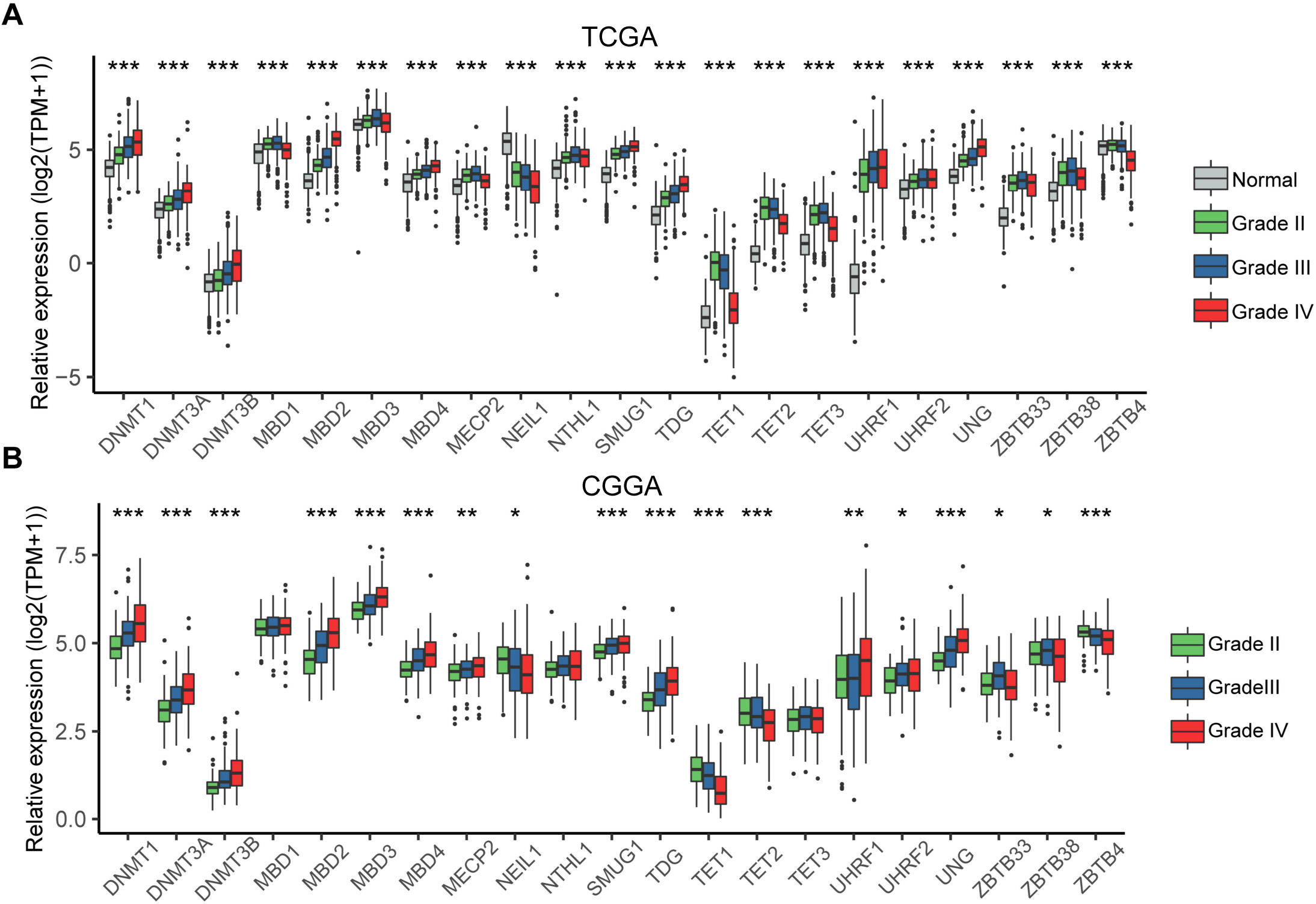
Overview of the 5mC regulators’ expression in glioma samples. The expression of 21 5mC regulator genes in glioma samples with different pathological grades in the TCGA (**A**) and CGGA (**B**) glioma cohorts. * indicates P < 0.05; ** indicates P < 0.01; *** indicates P < 0.001.

IDH mutation status is a well-established biomarker in glioma, and IDH wide-type usually indicates a relatively unfavorable prognosis (44). Next, we assessed the relationship between IDH status and the expression levels of 5mC regulators. In the TCGA dataset, the GBM subgroup comprised 147 IDH wide-type and 11 mutant samples, and the LGG subgroup consisted of 96 IDH wild-type and 423 mutant samples. To avoid a statistical bias results from the imbalanced numbers of samples, we only focused on the differences of 5mC regulators’ expression between IDH mutant and wild-type samples in the TCGA LGG cohort. As shown in Figure S1A, DNMT1, DNMT3A, MBD2, MBD3, MBD4, SMUG1, and UNG were highly expressed in the IDH wide-type group, while MBD1, TET1, TET2, TET3, and ZBTB4 were highly expressed in the IDH mutant group (P <0.05). In the CGGA cohort, 101 IDH wide-type and 42 mutant GBM samples, as well as 149 IDH wild-type and 175 mutant LGG samples, were included. We further investigated the overview of 5mC regulators’ expression to validate the previous findings in the CGGA cohort. We found that 50% of genes (6/12, DNMT3A, MBD2, MBD4, TET1, TET2, and TET3) in the CGGA dataset had a similar expression with those in the TCGA cohort (Figure S1B-C).

Combined deletion of both the short arm of chromosome 1 and of the long arm of chromosome 19 (1p19q co-deletion) is recognized as an indicator of favorable prognosis in gliomas, especially those with low grades (45). In LGG with IDH-mutation, 171 1p19q co-deleted and 252 non-codeleted samples were included in the TCGA cohort, while 57 1p19q co-deleted and 75 non-codeleted samples in CGGA cohort. Combining the results from two independent cohorts, we found that DNMT3A, MBD2, MBD3, UNG, and ZBTB33 were highly expressed in the co-deleted group (P <0.05), while the expression level of NEIL1 was much lower (P <0.01) (Figure S2).

We further investigated whether the expression changes of 5mC regulators in glioma were influenced by somatic mutation or copy number variation. 794 cases of gliomas with both the information of mutation and copy number alteration were enrolled, and we found that the frequencies of genetic alteration of 21 5mC regulators were much low in glioma, with less than 2% in each gene (Figure S3). This finding indicated that the differential expression patterns of these regulators in glioma did not originate from the genetic changes.

### Construction a novel glioma classification based on 5mC regulators

Considering the distinct expression patterns of 5mC regulators in gliomas with different tumor grades and molecular features, we further explored whether these genes can be used as a metagene to divide glioma into different subtypes efficiently. Gene expression data of these genes were obtained in the TCGA cohort and then subjected to the CNMF clustering algorithm. The average silhouette width was 0.94, 0.36, and 0.38 when samples were clustered into two, three, and four classes (Figure 2A and Figure S4). Combining with the clustering heatmap, these results demonstrated that CNMF achieved adequate robustness when all glioma samples were classified into two clusters, with cluster1 consists of 470 patients and cluster2 of 217 patients. PCA plot showed a clear distinction in gene expression profiles between these two subgroups (Figure 2B). We compared the difference of clinical outcomes between these two clusters, and we noticed that patients in cluster2 showed worse prognosis when compared with those of cluster1 (HR = 3.03, 95% CI = 2.26-4.08, P = 0) (Figure 2C).

**Figure 2.**
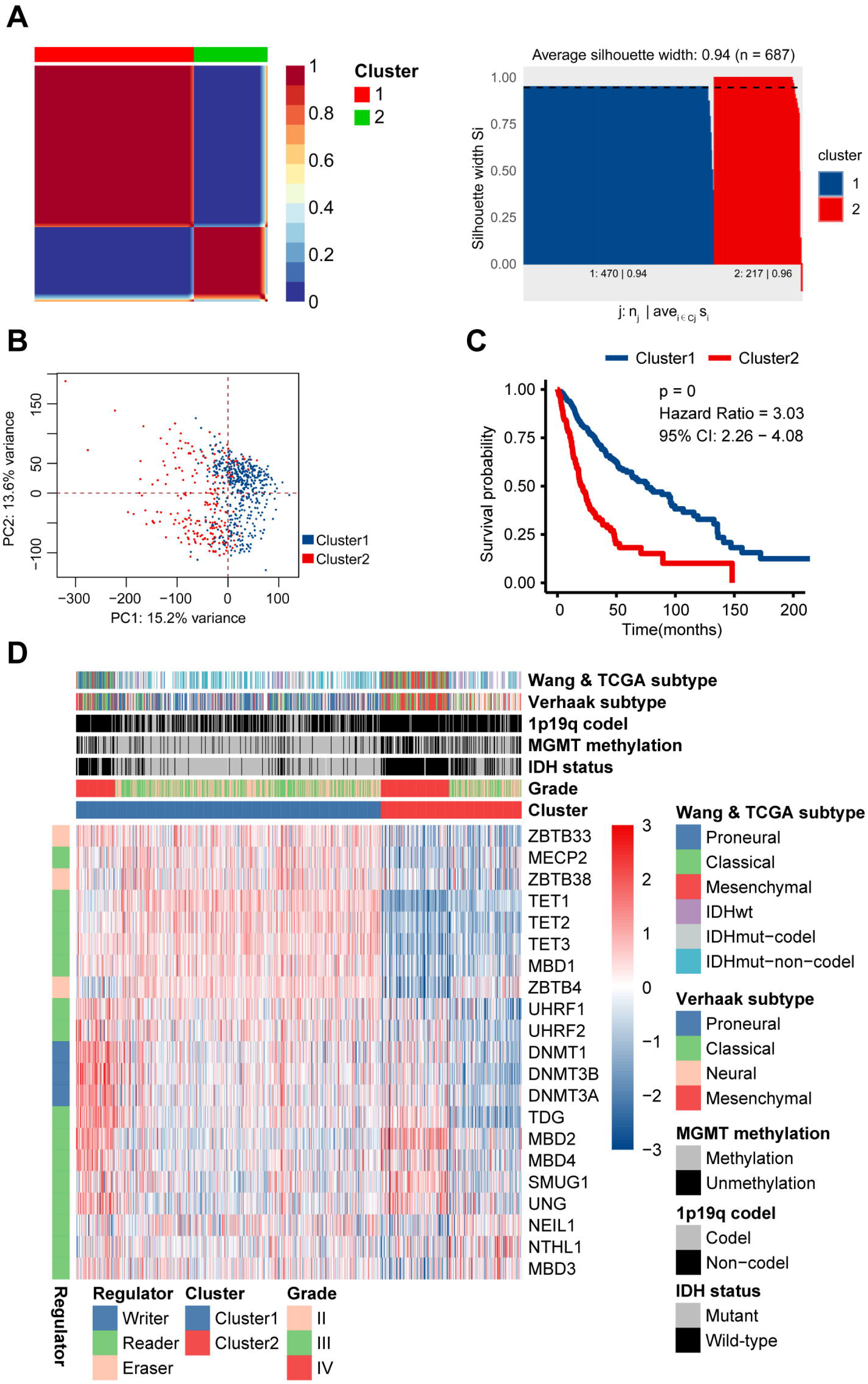
Identification of two distinct glioma subtypes using the CNMF method based on 5mC regulators. **A.** Heatmap of the sample similarity matrix and silhouette width plot of the subtypes when the number of subgroups was set as two. **B.** PCA (Principal Components Analysis) plot showing the difference in gene expression patterns between two clusters. The X-axis and Y-axis represent the top two principal components, respectively. **C.** Comparison of overall survival for patients of different clusters in the TCGA glioma cohort. **D.** Significant associations among the novel classification, clinicopathological characteristics, and well-established molecular subtypes.

The significant difference in prognosis between clusters reminded us that molecular might be the major reason. Here, we found that cluster2 had much higher fractions of older (P <0.001), GBM (48.39%, 105/217, P <0.001), IDH wild-type (69.48%, 148/213, P <0.001), MGMT unmethylated (46.46%, 92/198, P <0.001), 1p19q non-codel(91.55%, 195/213, P <0.001) patients (Figure 2D and Table S1). Interestingly, part of GBM samples was included in the cluster1 (36.36%, 60/165), while some LGG samples were also mingled in cluster2 (51.61%, 112/217). Normally, GBM represents a worse prognosis compared with LGG. However, the significant intratumoral heterogeneity of glioma stratifies the clinical outcomes. We investigated the associations between well-established molecular classifications and the novel subtypes we identified. Briefly, TCGA Verhaak classification classified glioma into Proneural, Neural, Classical, and Mesenchymal subtypes. Based on single-cell gene expression profiles, Wang 2017 GBM classification subgrouped GBM into Proneural, Classical, and Mesenchymal. Diffuse LGG were classified into IDH wild-type (IDHwt), IDH mutant with 1p19q codeletion (IDHmut-codel), and IDHmut-non-codel (IDH mutant with 1p19q non-codeletion) by TCGA group. Only 17 (28.33%, 17/60) and 8 (15.56%, 8/59) samples were mesenchymal in GBM samples of cluster1 according to the Verhaak and Wang classification, respectively. Furthermore, LGG samples of cluster2 comprising merely 13 (17.33%, 13/75) proneural and 7 (12.73%, 7/55) IDHmut-codel samples. These findings suggested that GBM or LGG samples in one cluster shared fewer similarities with their counterparts in another subtype in genetic level, and confirmed the feasibility of the 5mC regulator-based glioma classification.

### Association of identified classification with malignant signaling pathways in glioma

The significant differences in prognosis, molecular subtypes, as well as clinicopathological features between these 5mC regulator-based subgroups indicated that remarkably different signaling pathways might exist among them. We next assessed global gene expression changes in cluster2 relative to cluster1 samples by GSEA (Table S2). The results manifested a universal up-regulation of canonical tumor related signaling pathways, including epithelial–mesenchymal transition (NES = 2.43, FDR = 2.9e-4), interferon gamma response (NES = 2.32, FDR = 2.9e-4), coagulation (NES = 2.24, FDR = 2.9e-4), IL6 JAK STAT3 (NES = 2.18, FDR = 2.9e-4), TNFα via NFKB (NES = 2.12, FDR = 2.9e-4), angiogenesis (NES = 2.02, FDR = 2.9e-4), and hypoxia pathways (NES = 1.92, FDR = 2.9e-4) in cluster2 (Figure 3A). Notably,immune response related pathway was also enriched in cluster2 samples (NES = 2.38, FDR = 2.9e-4), which suggested that distinct tumor immune characteristics might exist among two clusters. Biological processes concerning immune cells migration comprising granulocyte (NES = 2.46, FDR = 1.7e-3), neutrophil (NES = 2.43, FDR = 1.7e-3), T cell (NES = 2.28, FDR = 1.7e-3), leukocyte (NES = 2.24, FDR = 1.7e-3), and immune processes, including humoral (NES = 2.47, FDR = 1.7e-3), adaptive (NES = 2.13, FDR = 1.7e-3), and innate immune (NES = 2.10, FDR = 1.7e-3). We also found that mesenchymal subtype glioma gene set was positively enriched in cluster2 (NES = 2.74, FDR = 9.9e-4), while the proneural signature was negatively enriched (NES = −3.96, FDR = 0.015). Another gene sets about mesenchymal and proneural glioma further confirmed this finding, with NES equals to 2.73 (FDR = 3.1e-4) and −4.2 (FDR = 0.014), respectively. Invasiveness, which is associated with mesenchymal features, were also found positively enriched in cluster2 (NES = 2.42, FDR = 9.9e-4). The GO and KEGG enrichment analyses of genes upregulated in cluster2 samples (log2 Fold-Change >1) indicated that these genes were significantly related to immune cells activation, migration, and proliferation, as well as extracellular structure organization, which is consistent with the results obtained from GSEA (Figure 3B and Table S3).

**Figure 3.**
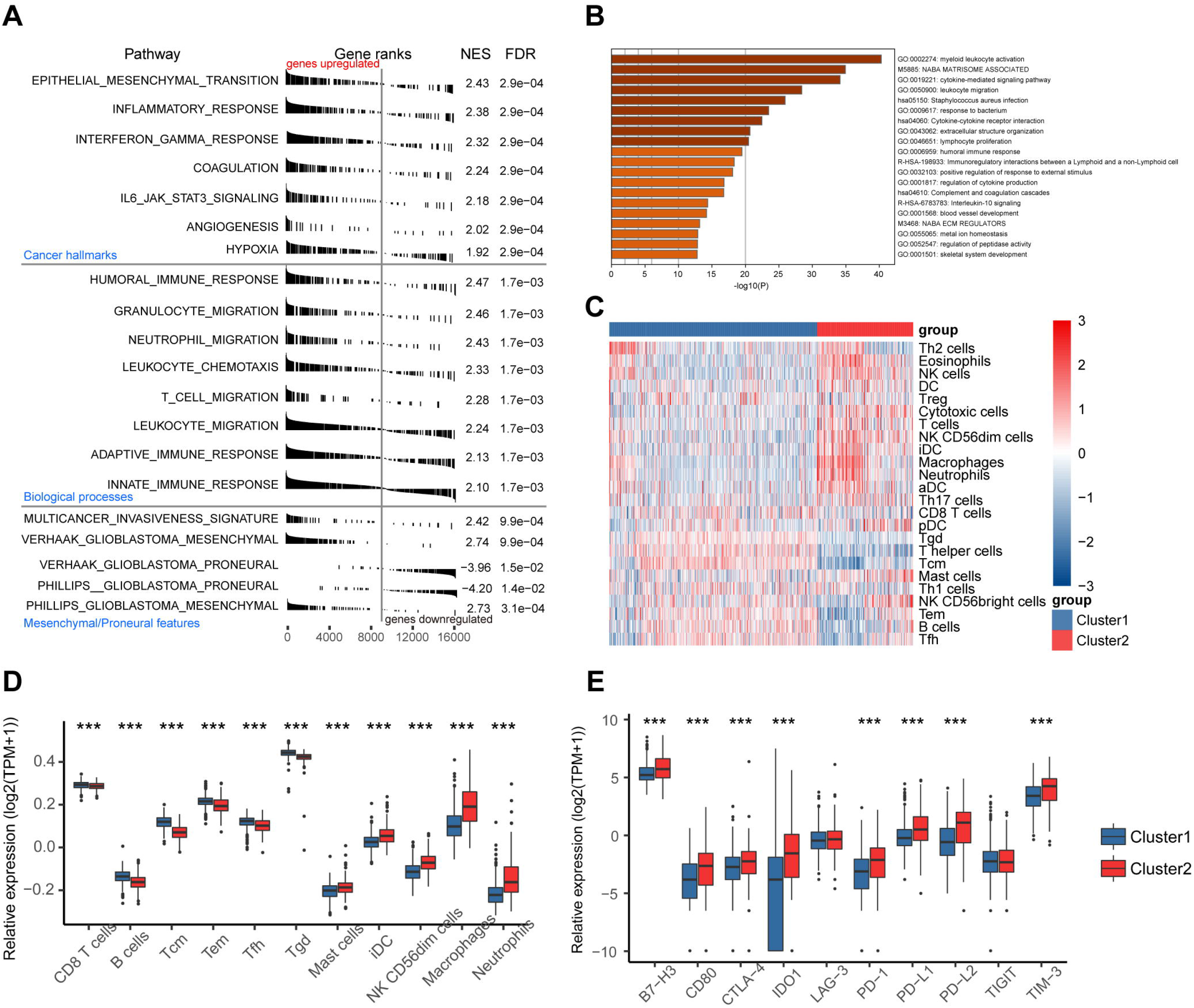
Functional annotation of the molecular differences and comparison of immunological features in different clusters. **A.** GSEA (Gene set enrichment analysis) showing the differences of underlying mechanisms between two clusters. Gene sets were divided into three subgroups, including pathways of cancer hallmarks, immune-related biological processes, and Proneural/Mesenchymal GBM specific genes. The label of “gene-upregulated” and “gene-downregulated” represent genes overexpressed/underexpressed in samples of cluster2 compared with cluster1. **B.** Enriched biological processes and signaling pathways in genes that were overexpressed in samples of cluster2. The X-axis indicates statistical significance. **C.** Comparison of the abundance of 24 types of infiltrated immune cells in glioma samples between different clusters. **D.** Boxplot visualizing significantly different immune cells between two clusters, including pro-tumor and anti-tumor immune cells. **E.** The expression levels of several immune checkpoints between two clusters. *** indicates P < 0.001.

The malignant microenvironment in solid tumors is a complex comprising both tumor cells and other non-tumor cells, including diverse immune cells. Immune cell infiltration in the two subtypes was estimated by ssGSEA, and the abundance of 24 kinds of immune cells was compared between cluster1 and cluster2 (Figure 3C). Higher contents of some pro-tumor immune cells were found in cluster2, such as mast cells, iDC, NK CD56dim cells, macrophages, and neutrophils (all P <0.001). Meanwhile, fewer fractions of anti-tumor immune cells were enriched in cluster2, including CD8 T cells, B cells, Tcm, Tem, Tfh, and Tgd (all P <0.001) (Figure 3D). We also assessed the expression levels of several well-known immune checkpoints in the TCGA cohort (B7-H3, CD80, CTLA-4, IDO1, LAG-3, PD-1, PD-L1, PD-L2, TIGIT, and TIM-3), and we found most of them were highly expressed in cluster2 (all P <0.001), except LAG-3 and TICIT (Figure 3E). Collectively, these results might explain the significant differences in prognosis and immune-related pathways between novel clusters.

### Identification of survival-related 5mC regulators and construction of risk model based on 5mC regulators

Similar expression patterns among 5mC regulators were observed, as mentioned above. Here, we further explore the potential interactions among these genes in glioma. By checking the gene list in the STRING database, we found several regulators may function as hub genes in regulating DNA methylation as they are the center nodes in the connection network, including DNMT3A, DNMT3B, TET2, MBD4, TDG, DNMT1, UHRF1 (Figure 4A). For example, the protein-protein interaction (PPI) network showed that the interactions between DNMT3B and UHRF1, TDG, DNMT3A, DNMT1, as well as UHRF2 were both experimentally confirmed and textmining supported. Meanwhile, the gene expression levels of DNMT3B and the other five potential closely associated genes were positively correlated (all spearman correlation coefficients (R) >0.5, P <0.05) in both TCGA and CGGA cohorts (Figure 4B). A similar scenario was observed for other hub genes in the network, suggested that these hub genes may function as key regulators in the development of glioma.

**Figure 4.**
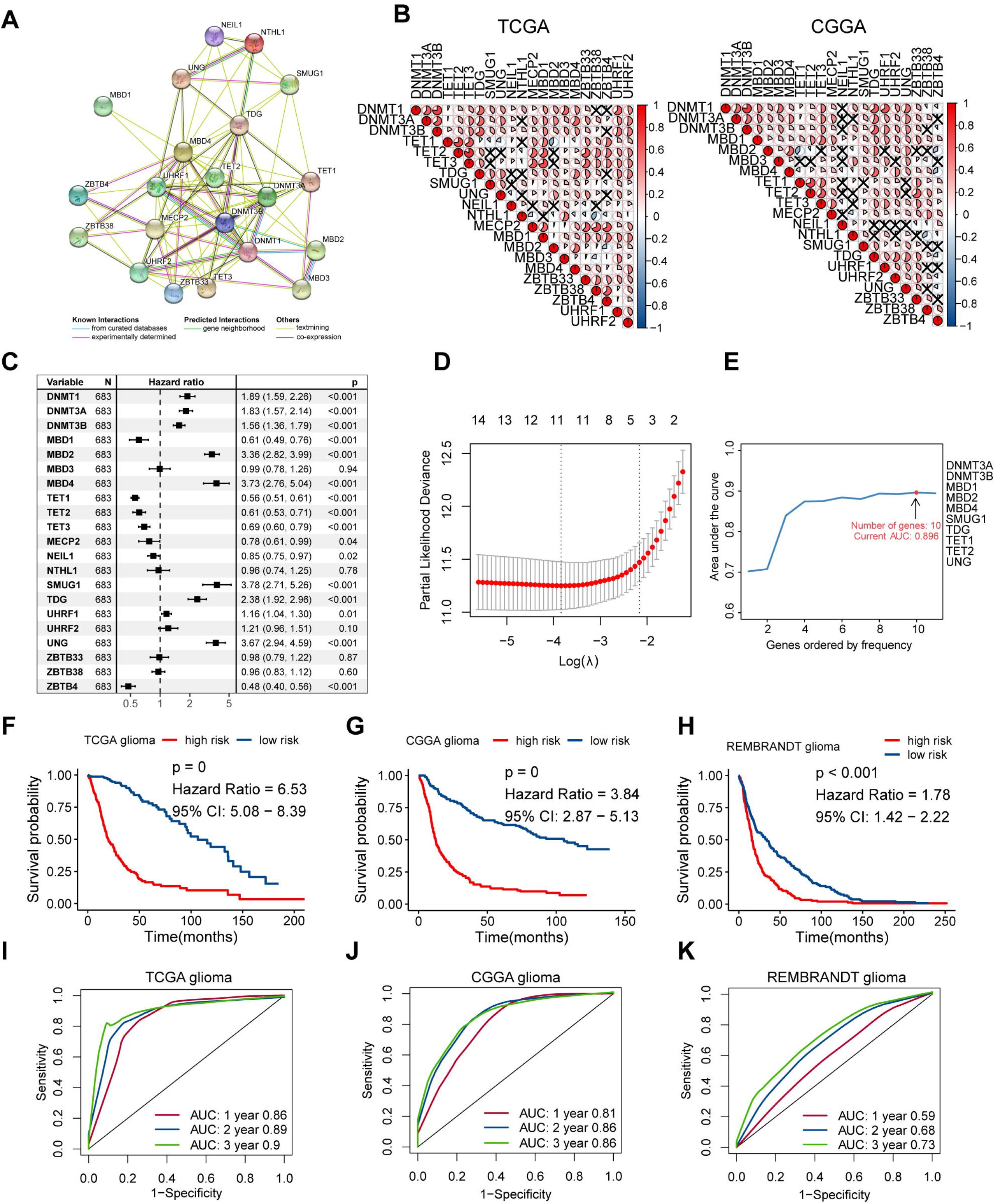
Construction of a prognostic risk model based on survival-related 5mC regulators in glioma. **A.** Protein-protein interaction (PPI) network of 21 5mC regulators. The label below showing the sources of these interactions. **B.** Correlations of the expression levels among 5mC regulators in both TCGA and CGGA cohorts. Each pie indicates the correlation coefficient (R) obtained from the Spearman correlation analysis, and the black cross represents no statistical significance. **C.** Identification of prognostic 5mC regulators in the TCGA cohort. **D.** Ten-fold cross-validation for selecting candidate genes in the LASSO model. The partial likelihood deviance was plotted using vertical lines with red dots, and the dotted vertical lines represent values based on minimum criteria and 1-SE criteria, respectively. **E.** Ten genes were adopted as optimal for survival prediction more than 500 times from 1000 iterations of lasso-penalized multivariate modeling. Comparison of overall survival for patients from high- and low-risk groups in the TCGA **(F)**, CGGA **(G)**, and REMBRANDT **(H)** glioma cohorts. Time-dependent ROC analysis for evaluating the performance of this gene signature in predicting 1-, 2- and 3-year survival of glioma patients in the TCGA **(I)**, CGGA **(J)**, and REMBRANDT **(K)** cohorts.

Sixteen survival-related 5mC regulators were identified using univariate Cox regression analysis in the TCGA cohort, and hub genes in the PPI network were all included in these prognostic factors (Figure 4C). Among these prognostic regulators, DNMT1, DNMT3A, DNMT3B, MBD2, MBD4, SMUG1, TDG, UHRF1, and UNG were risky genes with HR >1, while MBD1, TET1, TET2, TET3, MECP2, NEIL1, and ZBTB4 were favorable genes with HR <1. During LASSO regression analysis, the maximum prognostic value of the gene signature was confirmed when ten genes were enrolled in the signature (3-year area under the ROC curve (AUC) = 0.896, TCGA cohort), including DNMT3A, DNMT3B, MBD1, MBD2, MBD4, SMUG1, TDG, TET1, TET2, and UNG (Figure 4D-E). Subsequently, the risk score of each glioma patient was calculated by combining the expression values and corresponding coefficients of genes in the signature. Kaplan–Meier survival analysis showed that patients in the high-risk group had significantly worse prognosis compared with those with low risk in TCGA glioma cohort (HR = 6.53, 95% CI: 5.08-8.39, P = 0) (Figure 4F). The associations between risk score and clinicopathological factors and molecular features were illuminated in Figure S5. More samples with shorter survival time (P <0.0001), more frequencies of 1p19q non-codel (94.03% vs. 56.73%, P <0.0001), IDH wild-type (69.37% vs. 3.53%, P <0.0001), MGMT unmethylated (45.37% vs. 8.19%, P <0.0001), GBM (47.51% vs. 0.88%, P <0.0001), mesenchymal (36.08% vs. 0.37%, P <0.001), and cluster2 (49.56% vs. 14.04%, P <0.0001) samples were found in the high-risk group compared with those in low-risk group in TCGA cohort (Table S4).

We also validated the prognostic value of the 5mC regulator signature in different tumor grades as well as the different datasets. In CGGA and REMBRANDT glioma cohort, this signature also functioned as an unfavorable prognostic factor: HR = 3.84, 95% CI: 2.87-5.13, P = 0 in CGGA, and HR = 1.78, 95% CI: 1.42-2.22, P <0.001 in REMBRANDT (Figure 4G-H). Then, the TCGA glioma cohort was divided into GBM and LGG subgroup, and patients with high-risk still showed worse clinical outcomes than the low-risk group (Figure S6A, D). Similar results were found in CGGA-GBM, CGGA-LGG, REMBRANDT-GBM, and REMBRANDT-LGG cohort (Figure S6B, C, E, F). Time-dependent ROC curves were plotted to examine whether the risk model can serve as an effective index for predicting the prognosis of glioma patients at 1,2, and 3 years. The AUC was 0.86 at 1-year, 0.89 at 2-year, and 0.9 at 3-year in TCGA glioma cohort (Figure 4I). Meanwhile, we also tested the performance of the risk model in the CGGA and REMBRANDT glioma cohort (Figure 4J-K). Moreover, glioma patients in these three datasets were subgrouped according to the histology grade to further evaluate the clinical value of the 5mC regulator-based signature (Figure S7). Collectively, this signature was proved to behave as expected in predicting the prognosis for patients. Univariate and multivariate Cox regression analyses demonstrated that this gene signature could be used as an independent prognostic factor for diffuse glioma, GBM, and LGG patients both in TCGA and CGGA datasets (Table S5).

### High-risk GBM patients were more sensitive to temozolomide and radiotherapy

Temozolomide (TMZ) is the first-line chemotherapeutic drugs for high-grade gliomas, especially GBM (46). A combination of TMZ and radiotherapy can significantly prolong overall survival time for GBM patients. Here, we filtered the GBM subgroup from both the TCGA and CGGA cohorts and further carried out survival analyses for patients with high and low risk, respectively. In TCGA-GBM, patients without TMZ treatment showed significantly worse prognosis than those received TMZ treatment in the high-risk group (HR = 2.97, 95% CI: 1.67-5.26, P <0.001), while no difference was found between TMZ and non-TMZ treated patients in the low-risk group (P >0.05) (Figure 5A-B). We did not choose the CGGA-GBM cohort for validation because the detailed information of chemotherapeutic drug treatment was unavailable. As for radiotherapy, we found that the overall survival of patients without radiotherapy was much shorter than patients received radiotherapy in the high-risk group of TCGA-GBM (HR = 4.17, 95% CI: 1.82-9.51, P <0.001), while patients with low risk could not benefit from radiotherapy (P >0.05) (Figure 5C-D). Similar results were also found in the CGGA-GBM dataset (Figure 5E-F). These findings suggested that GBM patients with high risk scores may be more sensitive to TMZ treatment and radiotherapy, and therefore can benefit more from routine clinical treatment.

**Figure 5.**
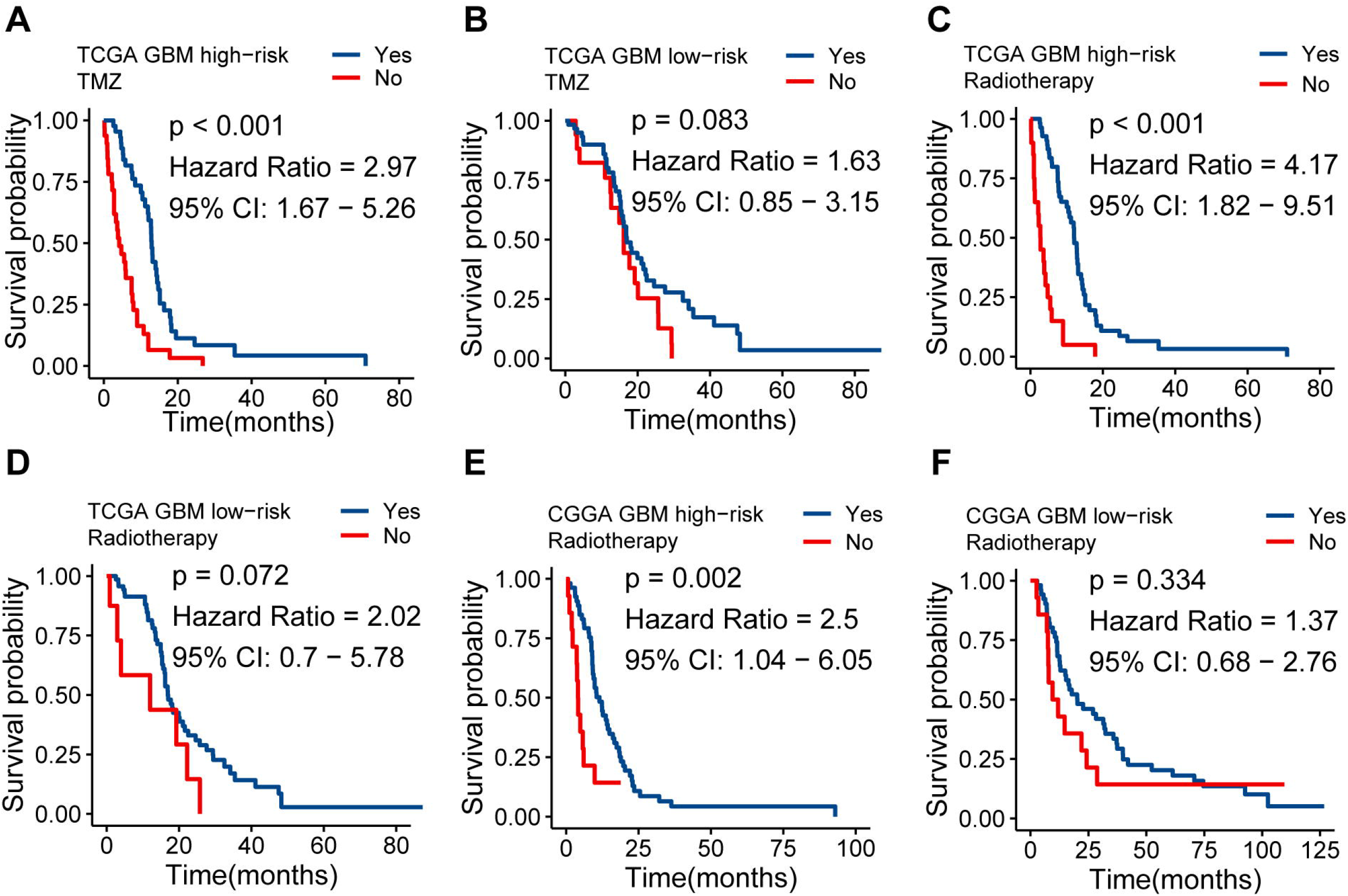
GBM patients with high risk scores were more sensitive to radiotherapy and TMZ treatment. Patients of the high-risk group can benefit more from TMZ-based chemotherapy **(A, B)** and radiotherapy **(C, D)** in the TCGA GBM cohort, compared with those of the low-risk group. A similar response to radiotherapy was found in the CGGA GBM cohort **(E, F)**.

### The 5mC regulator-based signature is closely related to tumor immunosuppression

As revealed above, 5mC regulator-based classification is associated with diverse immune responses and immune cell infiltration in glioma. More samples of cluster2 were found along with the increase of risk score, and this phenomenon indicated that 5mC regulator-based gene signature might be tightly related to anti-tumor immune processes. By estimating the abundance of infiltrated immune cells in glioma samples, we found that most cell types were statistically correlated with the increasing risk score both in TCGA and CGGA glioma cohorts (Figure 6A-B). Combining the results both from TCGA and CGGA datasets, pro-tumor immune cells were positively correlated with the risk score, such as Th2, NK CD56dim, iDC, macrophage, and neutrophil (all P <0.05), while the extent of infiltrated anti-tumor cells was inversely correlated with the risk score, including Tcm, Tem, Th1, NK CD56bright, Tfh, and Bcells (all P <0.05). Immune cells, as well as stromal cells in the tumor tissues, can regulate the tumorigenesis by numerous feedback machinery (47). We found that both immune and stromal scores calculated by the “ESTIMATE” method were positively related to risk score in TCGA and CGGA datasets (Figure 6A-B). Meanwhile, the purity of the solid tumor was decreasing along with the increasing risk score. Richard et al. defined a stroma-specific subtype of pancreatic ductal adenocarcinoma, and patients in the activated-stromal subtype had a worse survival time than those who belonged to the normal stromal subtype (42). Here, we further validated the relationship between risk signature and stromal cells, and the positive correlations were observed in the TCGA and CGGA cohort (Figure 6A-B).

**Figure 6.**
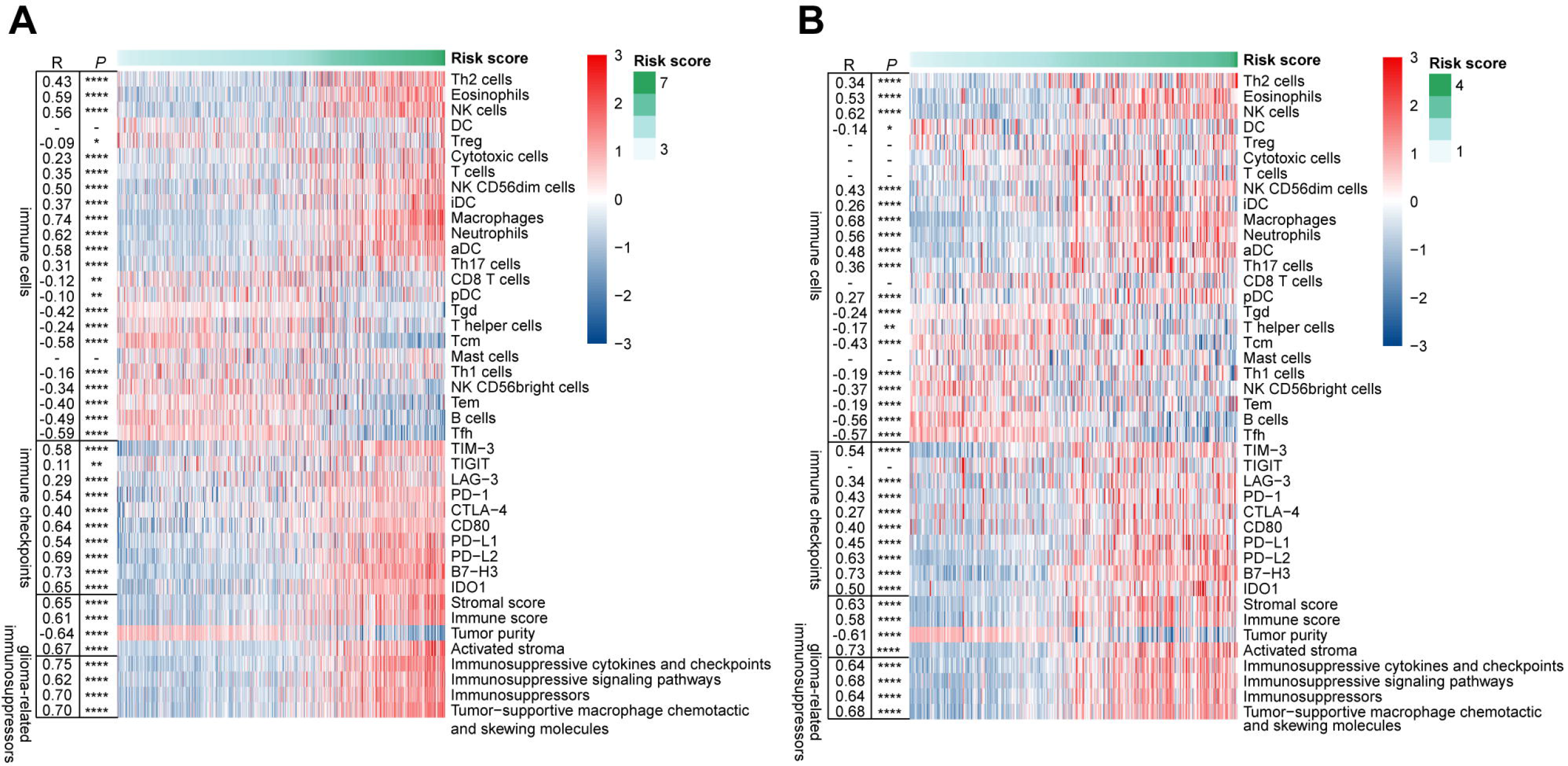
The gene signature is tightly associated with the intratumoral immune microenvironment in glioma. The relationships between risk score and immune cell infiltration, tumor purity, stroma abundance, immune checkpoints expression, as well as glioma-related immunosuppressive factors were estimated both in the TCGA **(A)** and CGGA **(B)** glioma cohorts. The correlation coefficients (R) were calculated by Pearson correlation analysis. * indicates P < 0.05; ** indicates P < 0.01; *** indicates P < 0.001.

Nine of ten immune checkpoint modules were positively correlated with increasing risk scores both in these two independent cohorts, except TIGIT, which indicated that the risk signature was implicated in immunosuppressive functions (Figure 6A-B). Significant positive relations were found between risk score and four glioma-related immunosuppressive factors both in TCGA and CGGA cohorts, including immunosuppressive cytokines and checkpoints, tumor-supportive macrophage chemotactic and skewing molecules, immunosuppressive signaling pathways, and immunosuppressors (all R >0.5, P <0.0001) (Figure 6A-B). These findings further solidify the association between 5mC regulator-based signature and tumor immune microenvironment and extend this mutual relationship to poor clinical outcomes.

### Establishment of a nomogram based on 5mC regulators

In order to develop a quantitative tool to predict the prognosis of glioma patients, we established a nomogram by integrating clinicopathological risk factors and 5mC regulator-based signature based on the multivariable Cox proportional hazards model (Figure 7A, C). The point scale of each row in the nomogram represents the point of each variable, and the survival likelihood of glioma patients was qualified by adding total scores of all variables together. The C-index of the nomogram reached 0.87 (95% CI: 0.86-0.89) and 0.78 (95% CI: 0.77-0.80) in TCGA and CGGA cohort, respectively. Calibration plots based on 1, 2, 3, and 5-year timepoints further confirmed the significant consistency between predicted and observed actual clinical outcomes of glioma patients in both two cohorts (Figure 7B, D). In summary, these findings confirmed that the nomogram was an optimal model for accurately predicting the clinical outcomes of glioma samples.

**Figure 7.**
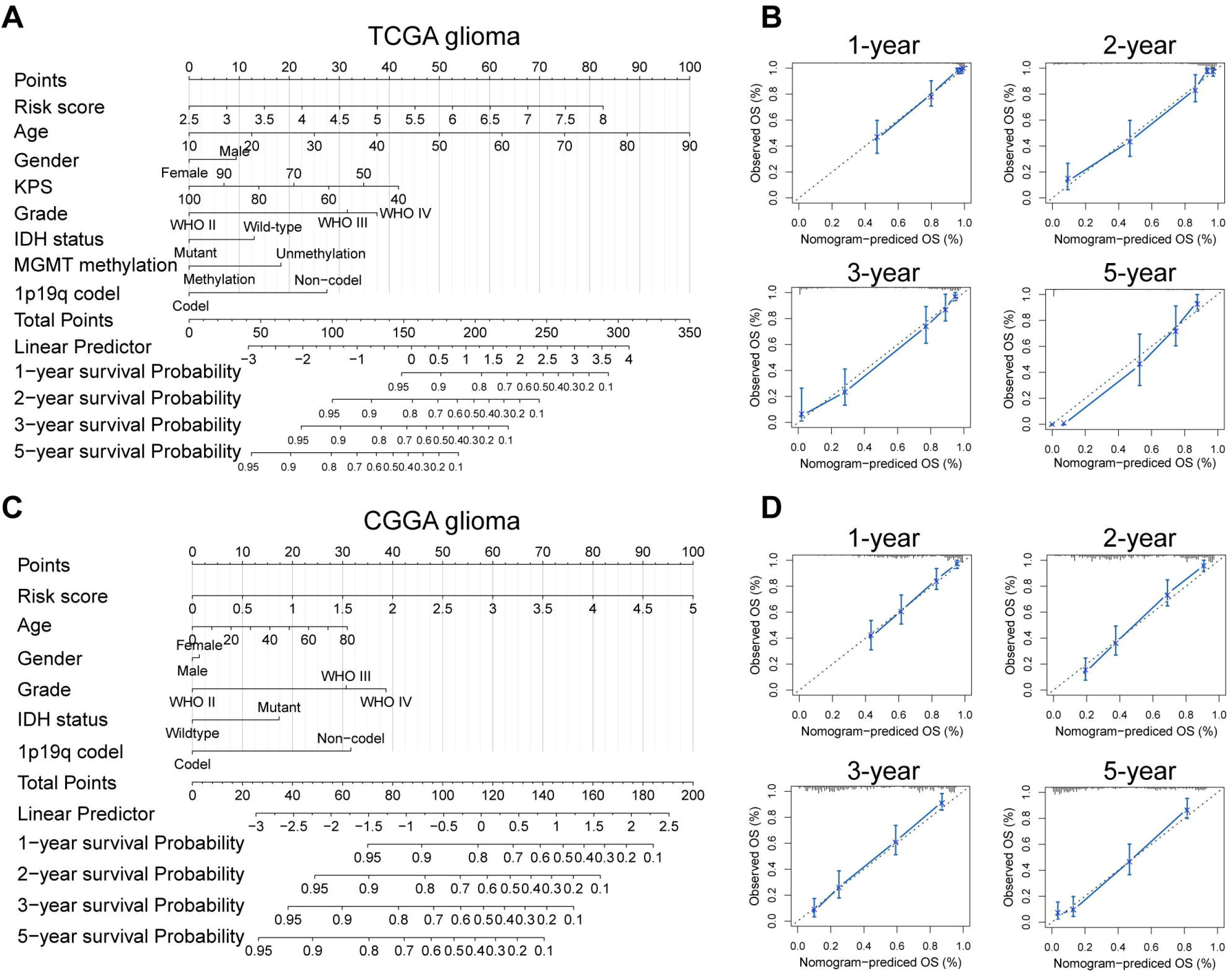
Developed nomogram to predict the probability of survival in glioma patients. Nomogram built with clinicopathological factors incorporated estimating 1-, 2-, 3-, and 5-year overall survival for glioma patients in the TCGA **(A)** and CGGA **(C)** cohorts. The calibration curves describing the consistency between predicted and observed overall survival at different time points in the TCGA **(B)** and CGGA **(D)** cohorts. The estimated survival was plotted on the X-axis, and the actual outcome was plotted on the Y-axis. The gray 45-degree dotted line represents an ideal calibration mode.

## Discussion

Abnormal methylation of CpG island in the promoter region of genes usually leads to transcriptional silencing of specific genes and therefore results in dysregulation of normal cellular biological processes (48). Toyota et al. (49) firstly discovered the CpG island methylator phenotype (CI MP) in colorectal tumor, which is defined as CpG island hypermethylation of genes enriched in a subgroup of tumor samples, and this conception has been widely investigated in various human cancers (50) (51). The previous study confirmed the existence of glioma-CpG island methylator phenotype (G-CIMP) and classified glioma into distinct subtypes based on G-CIMP features (52). The significant differences in histological grade, copy-number alterations, gene expression patterns, and prognosis between G-CIMP clusters highlights the importance of DNA-methylation in glioma. Here, we firstly investigated the expression of 5mC regulators in both TCGA and CGGA glioma cohorts. NEIL1 was less expressed in glioma samples compared with normal tissues, and negatively related with the increasing tumor grade. Any study on the expression of NEIL1 in glioma was not found by the literature search. The expression level of NEIL1 was inversely correlated with tumor mutation load (TML) in 13 types of human cancers in the TCGA database (R = −0.64, P = 0.019) (53). TML is associated with tumor grade in glioma, and high-grade gliomas demonstrate higher TML (54). Thus, we speculated that the predicted expression of NEIL1 in glioma is consistent with the actual outcome. G-CIMP is highly dependent on the mutation status of IDH, and almost all G-CIMP positive LGG samples harbor IDH mutation using unsupervised hierarchical clustering (55). Consistent with this, tight relationships between IDH mutation status and the expression levels of 5mC regulator genes were observed in our study.

Whole glioma samples from the TCGA cohort were stratified into two distinct subtypes based on the gene expression profiles of these 5mC regulators. This novel classification resulted in clear discrepancies in histological grade, clinical outcomes, and molecular features. However, not all GBM samples were included in cluster2, which is characterized as a worse prognosis. Previous studies have demonstrated that it is insufficient to classify glioma merely from the histological view (56). Verhaak et al. uncovered four subtypes of GBM using an 840 gene signature and revealed the heterogeneity in GBM (29). Moreover, Wang et al. modified this classification by single-cell sequencing as well as comprehensive validation, and deleted the Neural subtype since this kind of samples were always non-malignant. We linked our subtypes identified to these two well-established classifications and found that GBM samples in the cluster1 had a large proportion of non-mesenchymal, known as the less malignant phenotype. These findings made our novel classification more reasonable and feasible.

The discovery of multiple infiltrated immune cell types in CNS suggested that gliomagenesis is a biological process of immune disorder (57). Accumulating evidence shows that DNA methylation can regulate the transcriptomic landscape of the immune system, and dysregulation of this kind of epigenetic modification can result in immune-related diseases, including cancer (58). As critical regulatory factors, the role of 5mC regulatory genes in modulating tumor immune responses was further explored. Functional analysis identified significant upregulation of immune-related signaling pathways and immune responses, which indicated the potential mechanisms accounting for the significant differences between two subclasses. We quantified the abundance of infiltrated immune cells in solid tumor samples, and glad to see that the novel classification is closely associated with immune microenvironment remodeling. Anti-tumor immune cells, such as T cells, can be recruited into tumor loci and activated subsequently. Malignant cells can be protected from killing by these cells via immune escape machinery, including the PD-1/PD-L1 axis (59). Overexpressed immune checkpoint molecules in cluster2 further confirmed the vital role of 5mC regulators in regulating glioma-related immunological features.

Most studies merely focused on discovering gene signatures or biomarkers for GBM or LGG, while few attention has been paid on the whole gliomas. One of the main reasons is that the survival time differs a lot between different glioma grades: median overall survival of LGG patients is 6.9 years (60), while the counterpart of GBM is less than 15 months (61). The accuracy and sensitivity of those biomarkers will decrease when they were applied to another glioma cohort with a different pathological grade. Here, we built a risk model using ten 5mC regulator genes and confirmed its prognostic value in GBM, LGG, and whole glioma datasets. Furthermore, three independent glioma cohorts enrolled from different gene expression profiling platforms also determined the robustness and universality of this risk model. Among these genes in the signature, the specific functions of several genes have been intensively studied. For example, DNMT3A can reduce the sensitivity to TMZ treatment by interacting with ISGF3γ(62). The low expression level of TET1 will reduce the survival of GBM (63). Loss of the TET protein family causes the methylation of TIMP3, thus activates the apoptotic pathway and facilitates glioma growth (64). These findings may also be useful to develop novel therapeutic methods for glioma based on a single 5mC regulator or combined gene signature.

In conclusion, our study firstly analyzed the expression patterns and prognostic values of 5mC regulators in glioma. A novel molecular classification of glioma was successfully constructed, and the underlying role of intratumoral immune regulatory was identified. Furthermore, a 5mC regulator-based gene signature was developed and has been demonstrated to show satisfactory performance in predicting the clinical outcomes of glioma patients. Though further experiments should be carried out to clarify the cellular mechanisms of these crucial genes, this comprehensive study provides a novel scheme to facilitate our understanding of 5mC regulators in glioma.

## Notes

### Competing Interest Statement

The authors have declared no competing interest.

