## Supplementary material for "Molecular and clinical characterization of 5mC regulators in glioma: results of a multicenter study": Figure S

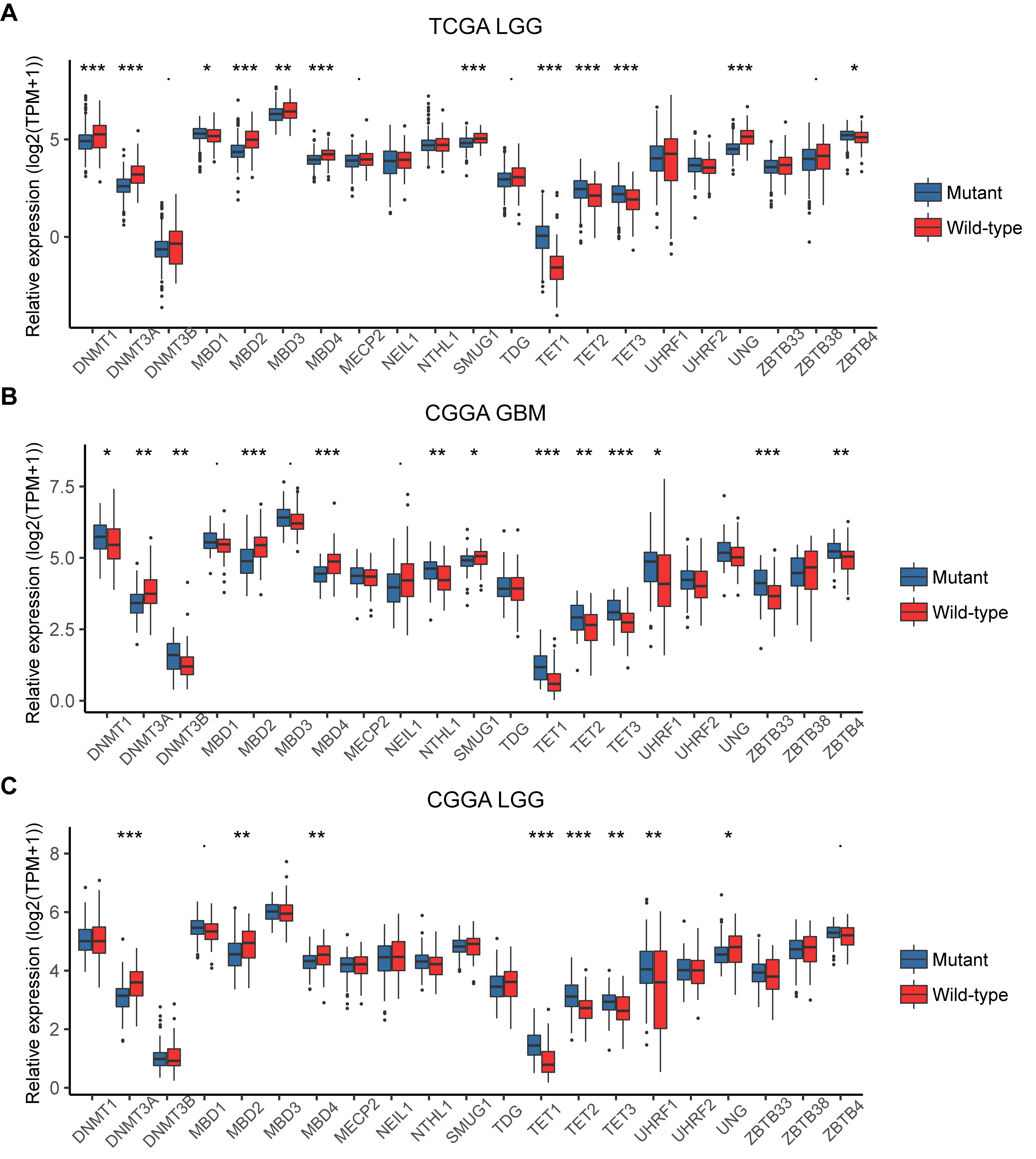

**Figure S1.** The associations between the expression of 5mC regulators and IDH mutation status. The expression levels of 5mC regulators were compared in IDH mutation and IDH wild-type samples in TCGA LGG **(A)**, CGGA GBM **(B)**, and CGGA LGG **(C)** cohorts. * indicates P < 0.05; ** indicates P < 0.01; *** indicates P < 0.001.

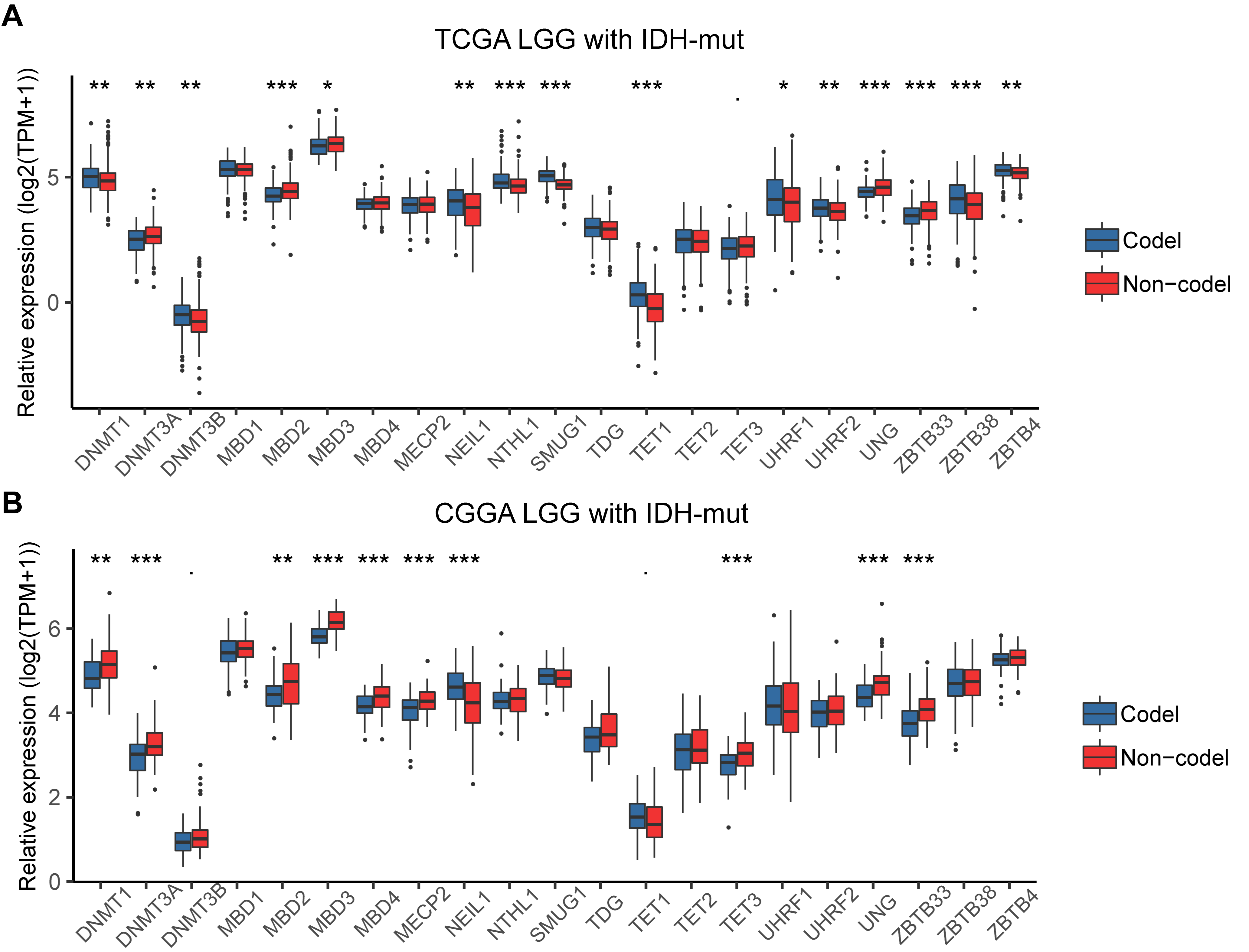

**Figure S2.** The associations between the expression of 5mC regulators and 1p/19q co-deletion status. The expression levels of 5mC regulators were compared in IDH mutant (IDH-mut) samples with 1p/19q codel and non-codel in TCGA LGG **(A)** and CGGA LGG **(B)** cohorts. * indicates P < 0.05; ** indicates P < 0.01; *** indicates P < 0.001.

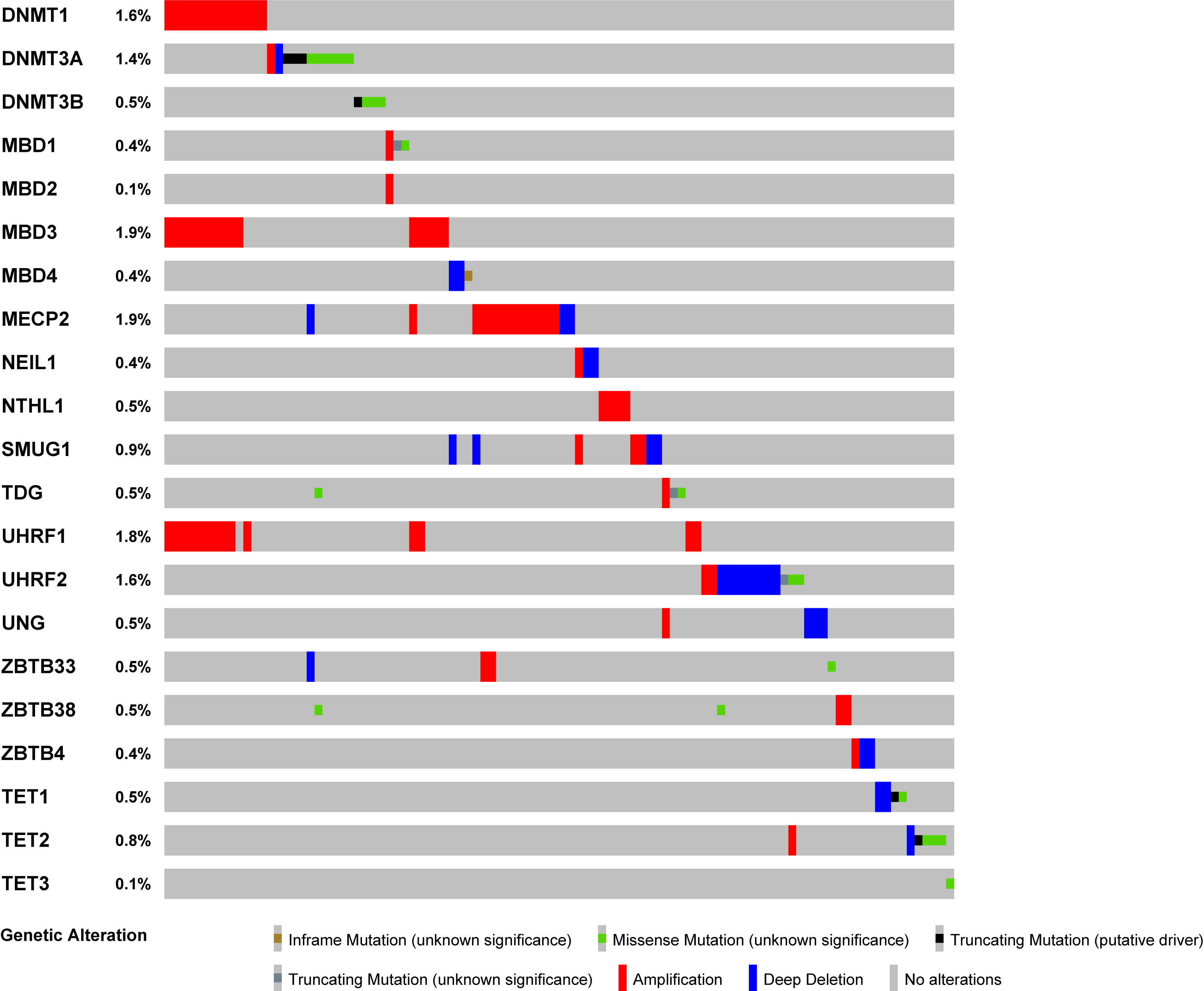

**Figure 3.** Genetic changes of 5mC regulators in the TCGA cohort. Overview of mutation and copy number variation of 5mC regulator genes in the TCGA glioma dataset.

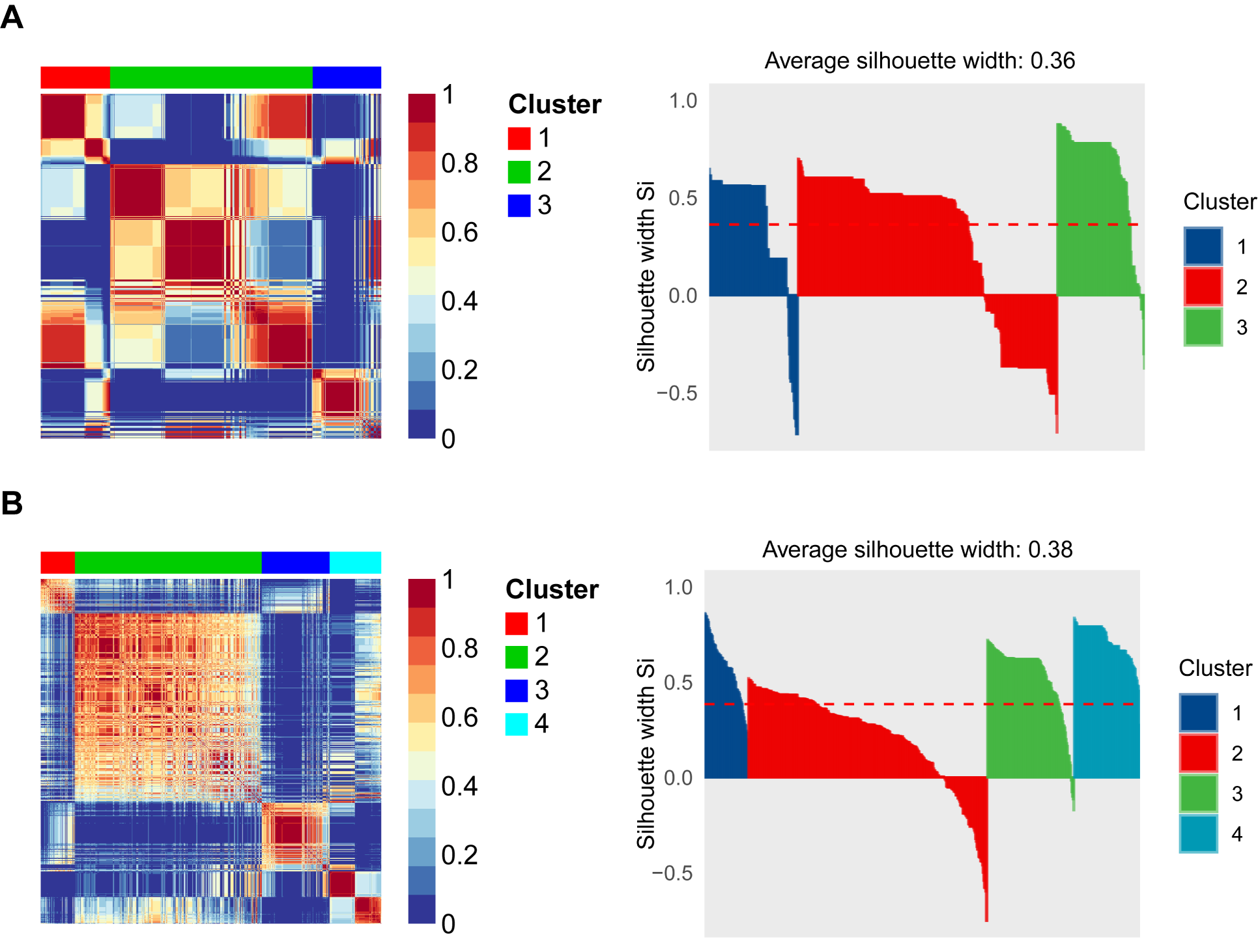

**Figure 4.** Clustering of whole glioma samples of the TCGA cohort using the CNMF method. Heatmaps of the sample similarity matrix and silhouette width plots of the subtypes when the number of subgroups was set as three **(A)** and four **(B)**.

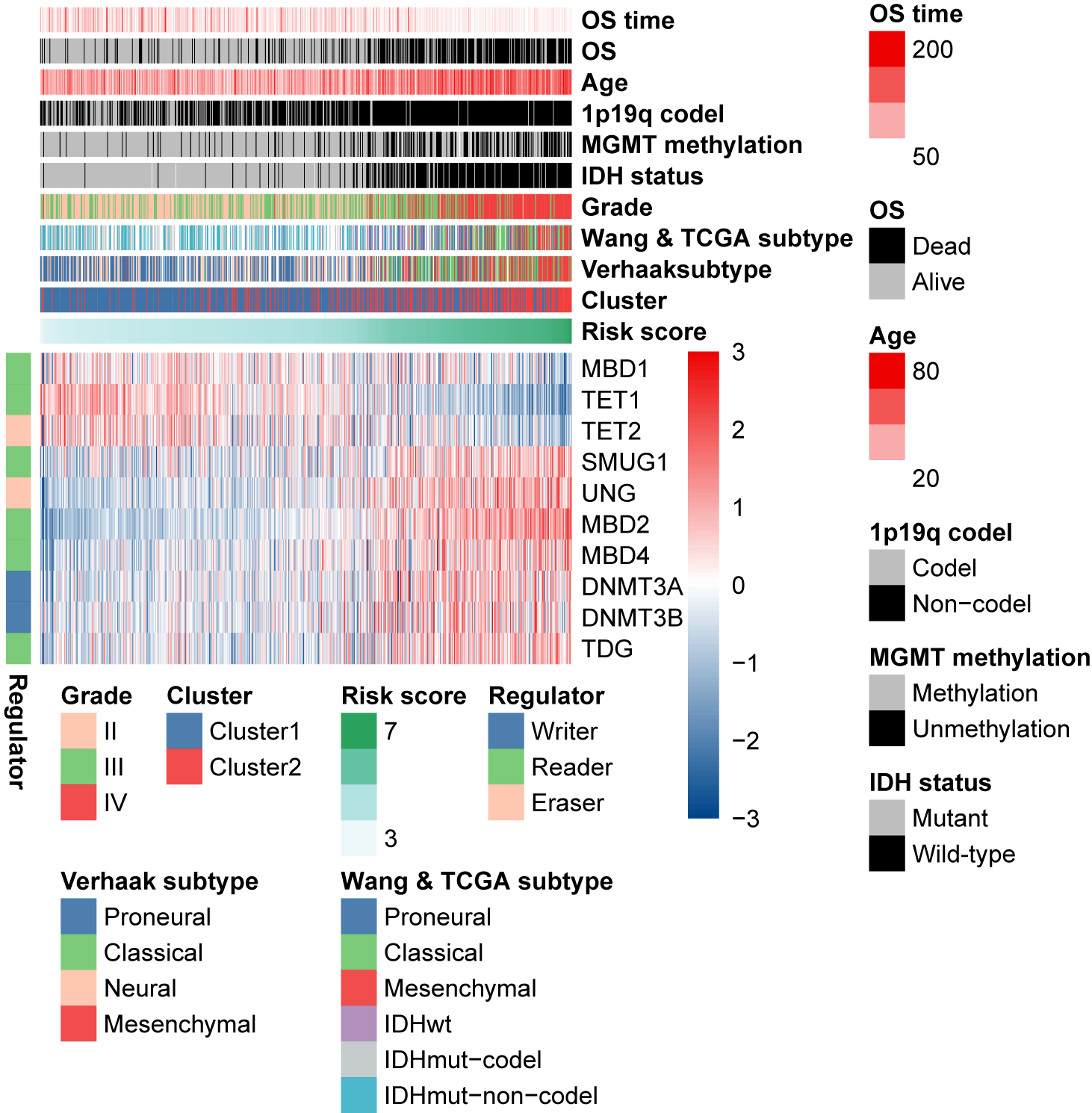

**Figure S5.** The relationship between risk score and clinicopathological features of glioma patients in the TCGA cohort. Heatmap shows the expression patterns of ten genes included in the signature, and the interactions between risk score and cluster, clinicopathological features, as well as well-acknowIedged molecular subtypes.

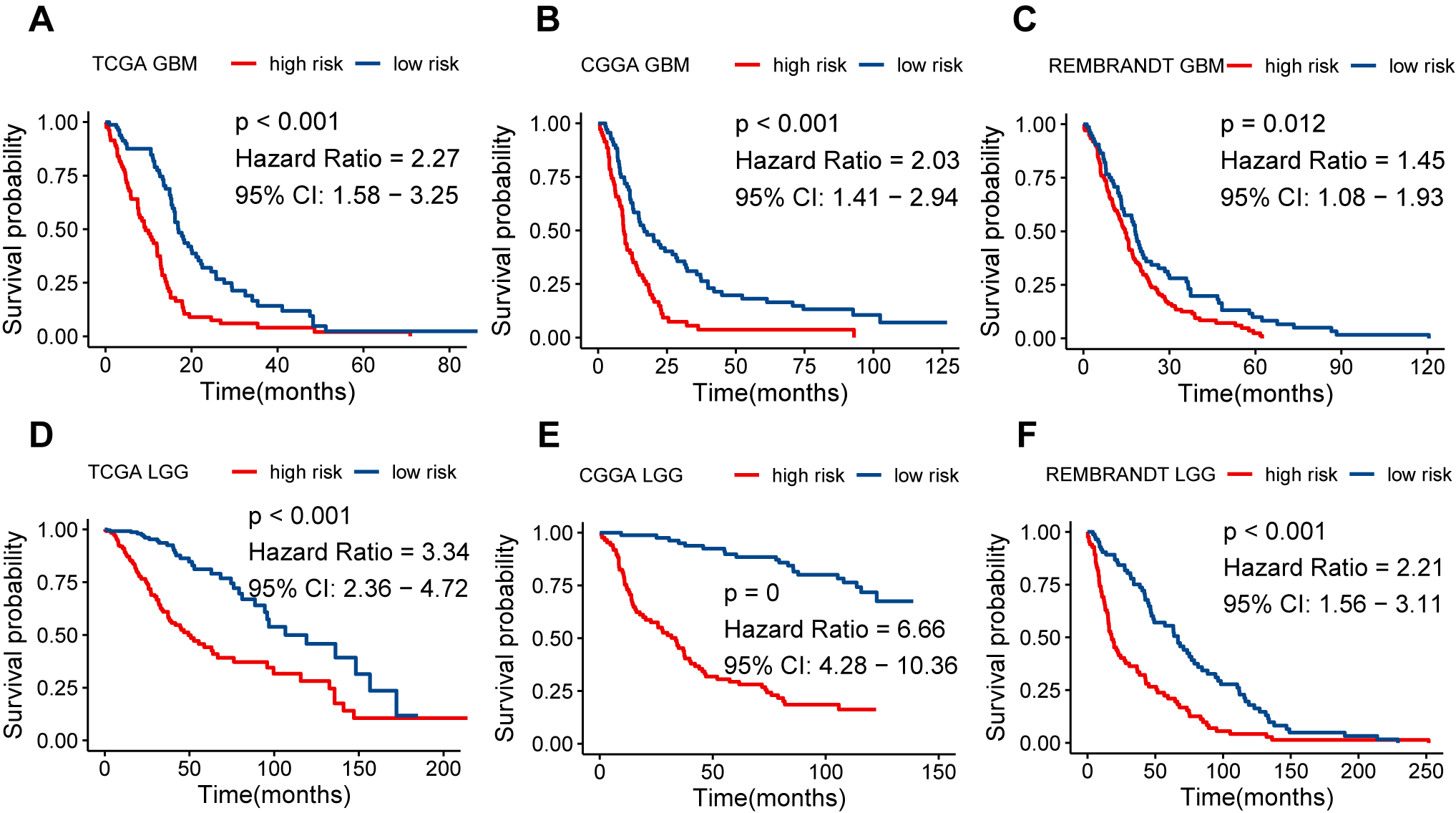

**Figure S6.** The prognostic value of the 5mC regulator-based risk model in GBM and LGG cohorts. Comparison of overall survival for both GBM **(A-C)** and LGG **(D-F)** patients from high- and low-risk groups in the TCGA, CGGA, and REMBRANDT cohorts.

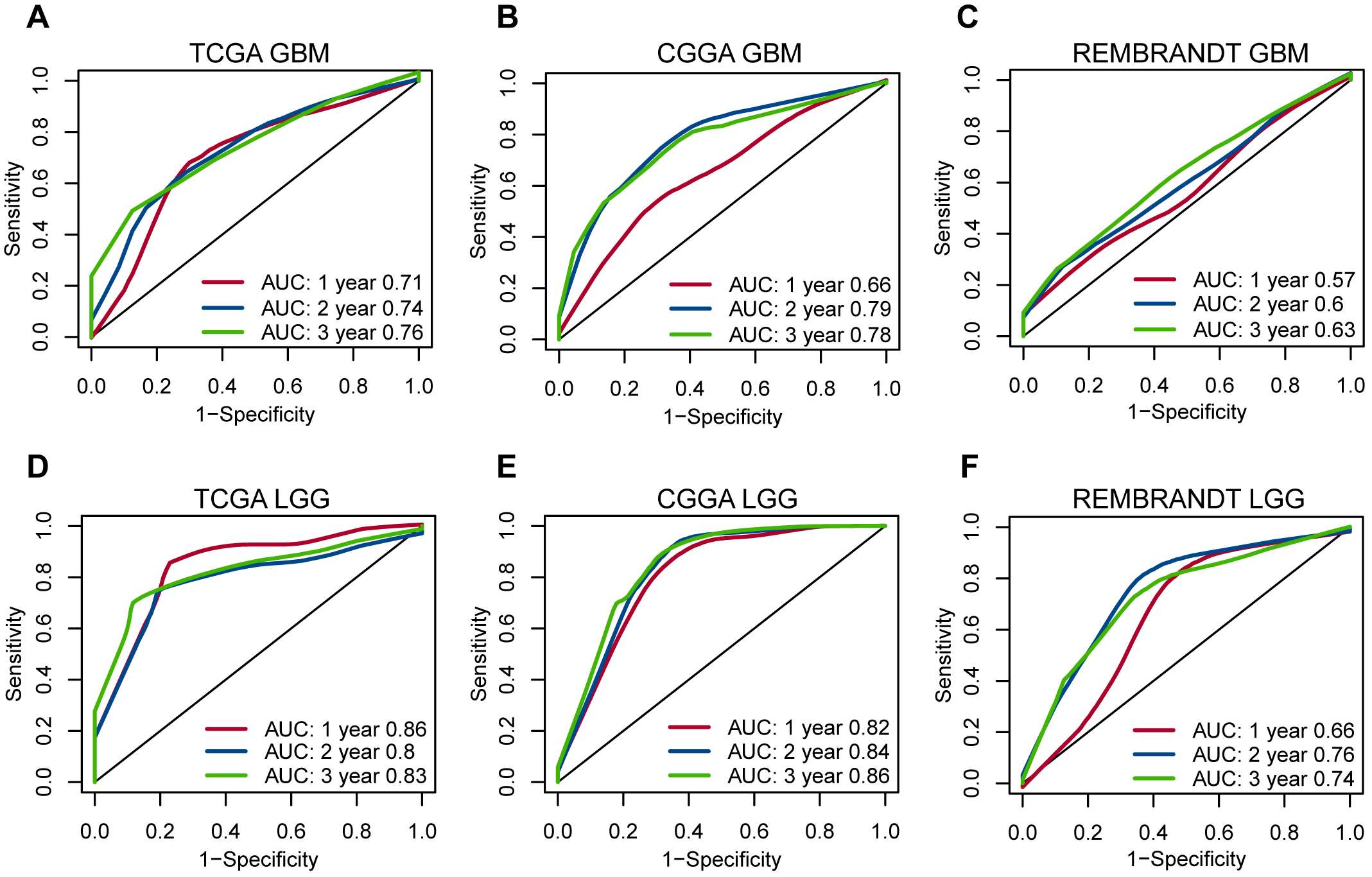

**Figure S7.** The 5mC regulator-based risk model is applicable for survival prediction for both GBM and LGG patients. Time-dependent ROC analysis for evaluating the performance of this gene signature in predicting 1-, 2- and 3-year survival of both GBM **(A-C)** and LGG **(D-F)** patients in the TCGA, CGGA, and REMBRANDT cohorts.
