## Supplementary material for "Molecular and clinical characterization of 5mC regulators in glioma: results of a multicenter study": Table S5

**Table S5.** Univariate and multivariate Cox regression of 5mC regulator-based risk model for overall survival in TCGA and CGGA glioma patients

TCGA glioma

| Variable | Univariate Cox Regression | | Multivariate Cox Regression | |
| --- | --- | --- | --- | --- |
|  | HR (95 % CI) | *P* | HR (95 % CI) | *P* |
| Age  >=60 *vs* <60 | 4.849 (3.731-6.3) | ***<0.001*** | 1.896 (1.263-2.846) | ***0.002*** |
| Gender  Male *vs* female | 1.222 (0.953-1.567) | 0.114 |  |  |
| KPS  >=80 *vs* <80 | 0.5 (0.344-0.725) | ***<0.001*** | 0.931 (0.607-1.428) | 0.744 |
| *IDH* status  Wild-type *vs* mutation | 9.347 (7.104-12.297) | ***<0.001*** | 2.049 (1.15-3.65) | ***0.015*** |
| *MGMT* methylation  Unmethylated *vs* methylated | 3.479 (2.673-4.528) | ***<0.001*** | 1.46 (0.976-2.186) | 0.066 |
| 1p19q status  Non-codel *vs* codel | 4.581 (2.956-7.1) | ***<0.001*** | 1.586 (0.773-3.251) | 0.208 |
| Risk score  High *vs* Low | 2.718 (2.428-3.044) | ***<0.001*** | 1.867 (1.506-2.315) | ***<0.001*** |

HR, hazards ratio; CI, confidence interval.

Note: Bold italics indicate statistically significant values (P < 0.05).

TCGA LGG

| Variable | Univariate Cox Regression | | Multivariate Cox Regression | |
| --- | --- | --- | --- | --- |
|  | HR (95 % CI) | *P* | HR (95 % CI) | *P* |
| Age  >=60 *vs* <60 | 5.021 (3.307-7.622) | ***<0.001*** | 3.42 (2.189-5.342) | ***<0.001*** |
| Gender  Male *vs* female | 1.116 (0.787-1.581) | 0.538 |  |  |
| KPS  >=80 *vs* <80 | 0.729 (0.361-1.472) | 0.378 |  |  |
| *IDH* status  Wild-type *vs* mutation | 6.042 (4.203-8.686) | ***<0.001*** | 1.656 (0.91-3.015) | 0.099 |
| *MGMT* methylation  Unmethylated *vs* methylated | 2.647 (1.818-3.854) | ***<0.001*** | 0.961 (0.606-1.525) | 0.866 |
| 1p19q status  Non-codel *vs* codel | 2.561 (1.617-4.058) | ***<0.001*** | 1.184 (0.703-1.995) | 0.526 |
| Risk score  High *vs* Low | 2.718 (2.269-3.256) | ***<0.001*** | 2.128 (1.621-2.794) | ***<0.001*** |

HR, hazards ratio; CI, confidence interval.

Note: Bold italics indicate statistically significant values (P < 0.05).

TCGA GBM

| Variable | Univariate Cox Regression | | Multivariate Cox Regression | |
| --- | --- | --- | --- | --- |
|  | HR (95 % CI) | *P* | HR (95 % CI) | *P* |
| Age  >=60 *vs* <60 | 1.323 (0.936-1.87) | 0.113 |  |  |
| Gender  Male *vs* female | 1.021 (0.713-1.463) | 0.909 |  |  |
| KPS  >=80 *vs* <80 | 0.824 (0.526-1.289) | 0.396 |  |  |
| *IDH* status  Wild-type *vs* mutation | 3.855 (1.672-8.887) | ***0.002*** | 3.164 (1.135-8.821) | ***0.028*** |
| *MGMT* methylation  Unmethylated *vs* methylated | 1.861 (1.23-2.815) | ***0.003*** | 1.644 (1.074-2.514) | ***0.022*** |
| Risk score  High *vs* Low | 2.718 (1.861-3.969) | ***<0.001*** | 2.319 (1.482-3.629) | ***<0.001*** |

HR, hazards ratio; CI, confidence interval.

Note: Bold italics indicate statistically significant values (P < 0.05).

CGGA glioma

| Variable | Univariate Cox Regression | | Multivariate Cox Regression | |
| --- | --- | --- | --- | --- |
|  | HR (95 % CI) | *P* | HR (95 % CI) | *P* |
| Age  >=60 *vs* <60 | 2.372 (1.614-3.487) | ***<0.001*** | 1.638 (1.084-2.475) | ***0.019*** |
| Gender  Male *vs* female | 0.931 (0.707-1.227) | 0.613 |  |  |
| *IDH* status  Wild-type *vs* mutation | 2.617 (1.98-3.458) | ***<0.001*** | 0.653 (0.441-0.966) | ***0.033*** |
| 1p19q status  Non-codel *vs* codel | 5.969 (3.612-9.862) | ***<0.001*** | 2.548 (1.462-4.44) | ***<0.001*** |
| chemotherapy  Yes *vs* no | 1.55 (1.154-2.082) | ***0.004*** | 0.994 (0.719-1.374) | 0.972 |
| Radiotherapy  Yes *vs* no | 0.519 (0.363-0.743) | ***<0.001*** | 0.673 (0.463-0.98) | ***0.039*** |
| Risk score  High *vs* Low | 2.718 (2.313-3.194) | ***<0.001*** | 2.589 (2.03-3.303) | ***<0.001*** |

HR, hazards ratio; CI, confidence interval.

Note: Bold italics indicate statistically significant values (P < 0.05).

CGGA LGG

| Variable | Univariate Cox Regression | | Multivariate Cox Regression | |
| --- | --- | --- | --- | --- |
|  | HR (95 % CI) | *P* | HR (95 % CI) | *P* |
| Age  >=60 *vs* <60 | 2.651 (1.366-5.147) | ***0.004*** | 0.835 (0.383-1.821) | 0.651 |
| Gender  Male *vs* female | 0.64 (0.421-0.972) | ***0.036*** | 0.725 (0.452-1.162) | 0.182 |
| *IDH* status  Wild-type *vs* mutation | 2.5 (1.605-3.895) | ***<0.001*** | 0.835 (0.49-1.423) | 0.507 |
| 1p19q status  Non-codel *vs* codel | 6.963 (3.582-13.533) | ***<0.001*** | 3.188 (1.505-6.754) | ***0.002*** |
| chemotherapy  Yes *vs* no | 2.152 (1.369-3.383) | ***<0.001*** | 1.601 (0.989-2.592) | 0.056 |
| Radiotherapy  Yes *vs* no | 0.527 (0.292-0.953) | ***0.034*** | 0.729 (0.382-1.39) | 0.337 |
| Risk score  High *vs* Low | 2.718 (2.163-3.416) | ***<0.001*** | 2.173 (1.61-2.932) | ***<0.001*** |

HR, hazards ratio; CI, confidence interval.

Note: Bold italics indicate statistically significant values (P < 0.05).

CGGA GBM

| Variable | Univariate Cox Regression | | Multivariate Cox Regression | |
| --- | --- | --- | --- | --- |
|  | HR (95 % CI) | *P* | HR (95 % CI) | *P* |
| Age  >=60 *vs* <60 | 1.598 (0.994-2.571) | 0.053 |  |  |
| Gender  Male *vs* female | 1.201 (0.826-1.748) | 0.337 |  |  |
| *IDH* status  Wild-type *vs* mutation | 1.028 (0.699-1.51) | 0.89 |  |  |
| chemotherapy  Yes *vs* no | 0.438 (0.294-0.651) | ***<0.001*** | 0.467 (0.311-0.701) | ***<0.001*** |
| Radiotherapy  Yes *vs* no | 0.631 (0.402-0.991) | ***0.046*** | 0.619 (0.392-0.977) | ***0.039*** |
| Risk score  High *vs* Low | 2.718 (1.828-4.042) | ***<0.001*** | 2.685 (1.728-4.17) | ***<0.001*** |

HR, hazards ratio; CI, confidence interval;

Note: Bold italics indicate statistically significant values (P < 0.05).
